# Matrix Viscoelasticity Regulates the Stemness and Multilineage differentiation of Primary Neural Progenitor-Stem Cells in 3D

**DOI:** 10.64898/2026.09.11.750834

**Authors:** Supeng Ding, Joo Ho Kim, Mengke Wang, Luo Gu

**Affiliations:** Department of Materials Science and Engineering, Johns Hopkins University, 3400 N Charles St, Baltimore, Maryland 21218, USA; Institute for NanoBioTechnology, Johns Hopkins University, 3400 N Charles St, Baltimore, Maryland 21218, USA; Translational Therapeutics and Regenerative Engineering Center, Johns Hopkins University School of Medicine, 400 N. Broadway, Baltimore, MD, 21231, USA

**Keywords:** stress relaxation, viscoelasticity, neural progenitor-stem cells, stemness, multilineage differentiation, mechanotransduction

## Abstract

Neural progenitor-stem cells (NPSCs) reside in mechanically dynamic brain microenvironments and give rise to neurons, astrocytes, and oligodendrocytes. Although recent studies have shown that matrix viscoelasticity can influence neural maturation and neurogenic differentiation, its role in primary NPSCs beyond neurogenesis remains less defined. How matrix viscoelasticity or stress relaxation regulates primary NPSC stemness, neuronal and glial differentiation, and the associated matrix-cell mechanotransduction pathways are not well understood. Here, we use alginate hydrogels with independently tunable stiffness and stress relaxation properties to investigate how matrix stress relaxation regulates the fate of primary subventricular zone (SVZ)-derived NPSCs in 3D. The results suggested that matrices with faster stress relaxation enhances stemness maintenance, radial glial-like marker expression, and differentiation of NPSCs toward neuronal, astrocytic, and oligodendrocytic lineages in the corresponding biochemical environments. In mixed neuronal/astrocytic differentiation conditions, fast-relaxing matrices preferentially promote neuronal differentiation. Mechanistically, NPSC responses to matrix stress relaxation involve integrin-mediated adhesion, actomyosin contractility, actin polymerization, and Piezo1 activity, with distinct contributions across differentiation lineages. Together, these findings reveal how matrix stress relaxation regulates primary NPSC stemness and multilineage differentiation through multiple mechanotransduction pathways in 3D.

## Main Text Introduction

Neural progenitor-stem cells (NPSCs) can self-renew and differentiate into neurons, astrocytes, and oligodendrocytes. The subventricular zone (SVZ) and subgranular zone (SGZ) are two well-studied NPSC niches in the brain (1). Because brain tissue is soft and viscoelastic, neural cells, including NPSCs, reside within and respond to this characteristic biophysical environment in vivo (2). Brain mechanical properties also change during aging and disease progression (3, 4), highlighting the importance of understanding how matrix mechanics influence NPSC behavior.

Traditional two-dimensional (2D) culture systems have been used to study how substrate stiffness regulates NPSC differentiation. Most studies reported enhanced neuronal differentiation on softer substrates (5–10), whereas astrocytic differentiation varies depending on culture medium and differentiation conditions (5–8, 11). However, 2D culture lacks essential three-dimensional (3D) cues and provides only a limited approximation of the natural cell microenvironment. The same mechanical cue can also produce different, or even opposite, outcomes in 2D and 3D culture. For example, fast stress relaxation has been reported to suppress NPSC neurogenesis on 2D substrates but promote neurogenesis in 3D matrices (12, 13).

Hydrogels are widely used to study cell behavior in 3D since. encapsulated cells experience matrix cues such as elasticity, viscoelasticity, porosity, etc. similar to physiological conditions. Previous studies have shown that matrix stiffness regulates NPSC proliferation and differentiation in 3D culture (14–16). In addition to stiffness, extracellular matrix (ECM) remodeling is another important aspect of the 3D mechanical environment, occurring either through biochemical remodeling by ECM proteins and enzymes (17–19), or through physical reorganization as cells deform the surrounding matrix (20). Highly degradable ECM-mimetic materials support NPSC stemness maintenance and differentiation capacity in 3D hydrogels (17, 19). These findings show that NPSCs are regulated not only by static stiffness, but also by dynamic matrix mechanical properties.

Among the dynamic matrix properties relevant to 3D neural microenvironments, matrix viscoelasticity is particularly important because it allows cells to physically remodel their surroundings over time rather than only sense a static stiffness. Prior studies have shown that this dynamic cue regulates several aspects of neural stem/progenitor cell behavior. In neurogenic contexts, highly viscoelastic hydrogels promoted neural maturation and neurite extension. This effect was mediated by actin polymerization-dependent mechanisms (21). Matrix stress relaxation also regulated neurogenesis in a dimensionality-dependent manner: compared with elastic conditions, viscoelastic matrices suppressed neurogenesis in 2D culture but promoted it in 3D, with spectrin mediating the 3D response (12, 13). In stemness and progenitor-organization contexts, viscoelastic N-cadherin-like matrix interactions supported neural progenitor cell stemness within 3D matrices (22). Stress relaxation timescale and hydrogel network connectivity regulated NPC stemness maintenance, proliferation, spatial organization, and differentiation capacity in engineered ELP-PEG hydrogels (23). Together, these studies establish matrix viscoelasticity as a regulator of neural stem/progenitor cell behavior in stemness maintenance, neural maturation, neurogenesis, and selected glial outcomes. However, because these findings were generated using different cell models, material systems, bioactive matrix contexts, and mechanical ranges with selected lineage outcomes, they do not yet define whether 3D matrix stress relaxation broadly coordinates primary NPSC stemness and multilineage fate potential within a single controlled platform.

This fragmentation leaves several gaps. First, it remains unclear whether matrix stress relaxation acts as a broad regulator of NPSC fate potential or whether its effects are restricted to specific stemness, neuronal, or glial contexts. A systematic investigation is therefore needed to evaluate stemness and neuronal, astrocytic, and oligodendrocytic differentiation within the same material platform and matched mechanical conditions.

Second, while human cell lines and pluripotent stem cell-derived neural models are valuable, primary SVZ-derived NPSCs provide an important model for studying progenitors from an endogenous neurogenic niche. SVZ-derived progenitors are associated with radial glial-like states marked by nestin, GFAP, and RC2 expression (24, 25), reflecting a complex neural identity in which stem/progenitor features overlap with glial-associated marker expression.

Third, because previously reported viscoelastic hydrogel systems often incorporate bioactive material components or engineered adhesive interactions, interpretation of stress relaxation effects can be complicated by material-derived biochemical cues. A comparatively simple matrix system is therefore needed to better examine how stress relaxation regulates primary NPSC fate.

Finally, because stress relaxation is a matrix-derived mechanical cue, its effects on NPSC fate require analysis of how mechanical information is transmitted from matrix to cells. Although prior studies identified mechanisms involving actin polymerization, spectrin, N-cadherin-like interactions, and β-catenin signaling (12, 13, 21–23), how adhesion-mediated matrix engagement, cytoskeletal contractility and remodeling, and mechanosensitive ion-channel activity contribute to stress-relaxation-mediated NPSC fate regulation remains less clear. This is especially relevant for neural progenitor and neural lineage cells, where mechanosensitive channels such as Piezo1 can couple extracellular mechanical cues to calcium-mediated intracellular signaling and neural stem/progenitor cell fate regulation (26–28).

In this work, we use alginate hydrogels with independently tunable stiffness and stress relaxation properties to investigate how matrix stress relaxation regulates the fate of primary NPSCs in 3D. We examine NPSC stemness maintenance, radial glial-like marker expression, differentiation toward neuronal, astrocytic, and oligodendrocytic lineages, and fate bias under mixed neuronal/astrocytic differentiation conditions. We further investigate the matrix-cell mechanotransduction pathways involved in these responses, including integrin-mediated adhesion, actomyosin contractility, actin polymerization, and Piezo1 activity. Together, this study reveals how matrix stress relaxation regulates primary NPSC stemness and differentiation lineages in 3D through multiple mechanotransduction pathways.

## 1. Results

### 1.1. 3D culture of SVZ-derived NPSCs in viscoelastic hydrogels restores radial glial-like marker expression

We used previously developed alginate hydrogels in which stiffness and stress relaxation can be independently tuned (29). Hydrogels were prepared at two stiffness levels, approximately 7 kPa and 15 kPa, and three stress relaxation conditions: slow, medium, and fast. Stress relaxation was characterized by the stress-relaxation half-time, t_1/2_, defined as the time required for the initial stress to relax to half of its original value under constant strain. The slow-, medium-, and fast-relaxing hydrogels exhibited t_1/2_ values of approximately 3600 s, 600 s, and 150 s, respectively (Fig. 1a-c). RGD (arginine-glycine-aspartate) peptides were covalently coupled to alginate to support cell adhesion.

**Figure 1.**
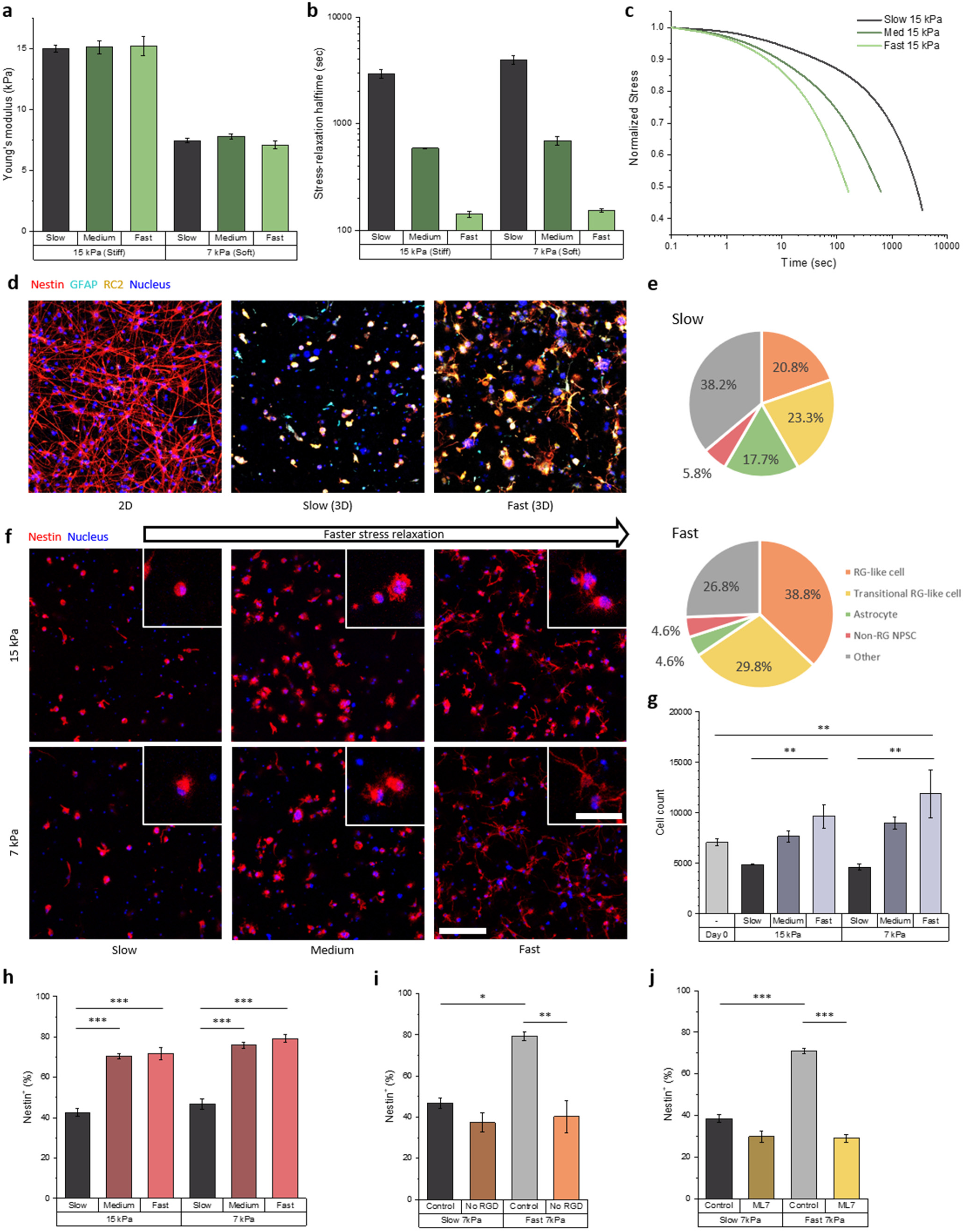
Fast matrix stress relaxation enhances radial glial-like marker expression and NPSC stemness maintenance. (a) Initial Young’s modulus of alginate hydrogels (N = 3). (b) Stress relaxation half-time of alginate hydrogels tuned by alginate molecular weight (N = 3). (c) Representative stress relaxation curves of slow-, medium-, and fast-relaxing hydrogels at 15 kPa under constant strain. (d) Immunocytochemistry showing nestin, GFAP, and RC2 expression in 2D culture and in 3D slow- and fast-relaxing hydrogels. Red, nestin; cyan, GFAP; yellow, RC2; blue, nuclei. (e) Quantification of nestin, GFAP, and RC2 marker-expression combinations in slow- and fast-relaxing 7 kPa hydrogels (N = 3). Radial glia-like (RG-like) cell: nestin^+^GFAP^+^RC2^+^; transitional RG-like cell: nestin^+^GFAP^−^RC2^+^; astrocyte: nestin^−^GFAP^+^RC2^−^; non-RG NPSC: nestin^+^GFAP^−^RC2^−^; Other: other combinations of nestin, GFAP, and RC2. (f) Representative images showing nestin expression and cell morphology in hydrogels with different stiffness and stress relaxation properties. Red, nestin; blue, nuclei. (g) Quantification of nuclei number per field of view after 7 days of culture compared with day 0. (h) Quantification of nestin- positive cell population (N = 3). (i) Quantification of nestin-positive cell population in slow- and fast-relaxing hydrogels with or without RGD ligands (N = 3). (j) Quantification of nestin-positive cell population in slow- and fast-relaxing hydrogels with or without ML7 treatment (N = 3). *p < 0.05, **p < 0.01, ***p < 0.001, by ANOVA with Tukey post-hoc test. Data are shown as means ± s.e.m. Scale bars: 150 µm; 50 µm in insets.

Primary NPSCs were isolated from the subventricular zone (SVZ) of neonatal rat brain lateral ventricles and expanded in two-dimensional (2D) culture in NPSC maintenance medium (30). The cells were then encapsulated in alginate hydrogels with different stiffness and stress relaxation properties and cultured in the same maintenance medium. In vivo, radial glial cells expressing nestin, GFAP, and RC2 give rise to SVZ-NPSCs, and postnatal SVZ-NPSCs can also express these markers (24, 31, 32). After 2D expansion, the NPSCs retained nestin expression but showed little or no GFAP and RC2 expression (Fig. 1d), consistent with prior reports that radial glial marker expression can change with developmental stage and in vitro culture conditions (33–35).

When 2D-expanded NPSCs were subsequently cultured in 3D viscoelastic alginate hydrogels for 7 days, GFAP and RC2 expression became detectable again, with more pronounced expression in fast-relaxing hydrogels (Fig. 1d). Among nestin-positive cells, the main populations were nestin^+^GFAP^+^RC2^+^ cells, consistent with a radial glial-like marker profile, and nestin^+^GFAP^−^RC2^+^ cells, which may represent an intermediate marker state between 2D-expanded NPSCs and radial glial-like progenitor cells (Fig. 1e) (32, 36). These marker-expression patterns indicate that 3D culture in viscoelastic environment supports restoration of *in vivo* radial glial-like progenitor features in SVZ-derived NPSCs, with fast stress relaxation condition showing a stronger effect.

Given the more pronounced radial glial-like marker expression of NPSCs in fast-relaxing hydrogels, we next examined whether matrix stress relaxation also influenced NPSC stemness maintenance and multilineage differentiation.

### 1.2. Fast matrix stress relaxation enhances NPSC stemness maintenance

To investigate how matrix stress relaxation regulates NPSC stemness maintenance, NPSCs were cultured in alginate hydrogels with two stiffness levels, approximately 7 kPa and 15 kPa, and three stress relaxation conditions: slow, medium, and fast. NPSCs were cultured in these hydrogels in NPSC maintenance medium for 7 days. Under both stiffness conditions, NPSCs showed greater spreading in faster-relaxing hydrogels (Fig. 1f). Cell number increased after 7 days only in fast-relaxing hydrogels, suggesting that faster stress relaxation supported NPSC survival and/or proliferation (Fig. 1g). Nestin expressing cell population among NPSCs was significantly lower in slow-relaxing hydrogels than in medium- and fast-relaxing hydrogels, indicating that NPSC stemness was better maintained in faster-relaxing matrices (Fig. 1h). This trend was observed at both 7 kPa and 15 kPa, showing consistent effect of matrix viscoelasticity at different stiffness (Fig. 1f-h).

We next examined whether integrin-mediated adhesion and actomyosin contractility contribute to stress-relaxation-dependent maintenance of NPSC stemness. To assess the role of integrin binding, NPSCs were cultured in slow- and fast-relaxing hydrogels with or without RGD ligands for 7 days. In fast-relaxing hydrogels, removal of RGD significantly reduced the percentage of nestin-positive cells to a level comparable to that observed in slow-relaxing hydrogels (Fig. 1i). In contrast, removing RGD did not significantly change nestin expression in slow-relaxing hydrogels.

To assess the role of actomyosin contractility, we inhibited myosin light chain kinase (MLCK) using ML7. MLCK inhibition similarly decreased the proportion of nestin-positive cells in fast-relaxing hydrogels to a level comparable to that in slow-relaxing hydrogels, while having no significant effect in slow-relaxing hydrogels (Fig. 1j). These results indicate that integrin-mediated adhesion and actomyosin contractility are required for matrix stress relaxation to regulate NPSC nestin expression. Removal of RGD ligands or inhibition of MLCK also reduced NPSC spreading in fast-relaxing hydrogels, although spreading remained greater than in slow-relaxing hydrogels (Fig. S1), indicating that the spreading response to fast stress relaxation is partly dependent on these pathways.

Because the quantity of calcium sulfate crosslinker was varied to tune hydrogel mechanics, we also examined whether the stemness outcome could be primarily explained by possible difference in initial Ca^2+^ crosslinker concentration. When the proportion of nestin-positive cells plotted against Ca^2+^ crosslinker concentration, the data did not show a consistent positive or negative relationship between Ca^2+^ concentration and NPSC stemness maintenance (Fig. S2). Instead, nestin expression was strongly associated with stress-relaxation condition, with higher nestin-positive population in medium- and fast-relaxing hydrogels than in slow-relaxing hydrogels. Within each stress-relaxation group, differences in initial Ca^2+^ crosslinker concentration corresponded to differences in gel stiffness; however, stiffness did not significantly affect nestin expression (Fig. 1h). These results support that the enhanced stemness maintenance in faster-relaxing hydrogels is associated with matrix stress relaxation rather than initial Ca^2+^ crosslinker concentration.

Together, these findings show that fast matrix stress relaxation enhances NPSC stemness maintenance, and that this response requires integrin-mediated adhesion and actomyosin contractility.

### 1.3. Fast matrix stress relaxation enhances neuronal differentiation of NPSCs

To investigate how matrix stress relaxation regulates neuronal differentiation, NPSCs were cultured in the same set of alginate hydrogels and induced toward neuronal differentiation using a two-step protocol (37). After differentiation, NPSCs cultured in fast-relaxing hydrogels showed higher proportion of cells positive of neuronal marker TUBB3 (β III tubulin) than those cultured in slow- or medium-relaxing hydrogels (Fig. 2a, b). Fast-relaxing hydrogels also increased the cell population that expresses the mature neuronal marker MAP2ab (Fig. 2c and Fig. S3). This trend was observed at both stiffness levels, whereas stiffness within the tested range did not significantly affect TUBB3 or MAP2ab expression (Fig. 2b, c). These results indicate that faster matrix stress relaxation enhances neuronal differentiation of NPSCs under neurogenic differentiation conditions.

**Figure 2.**
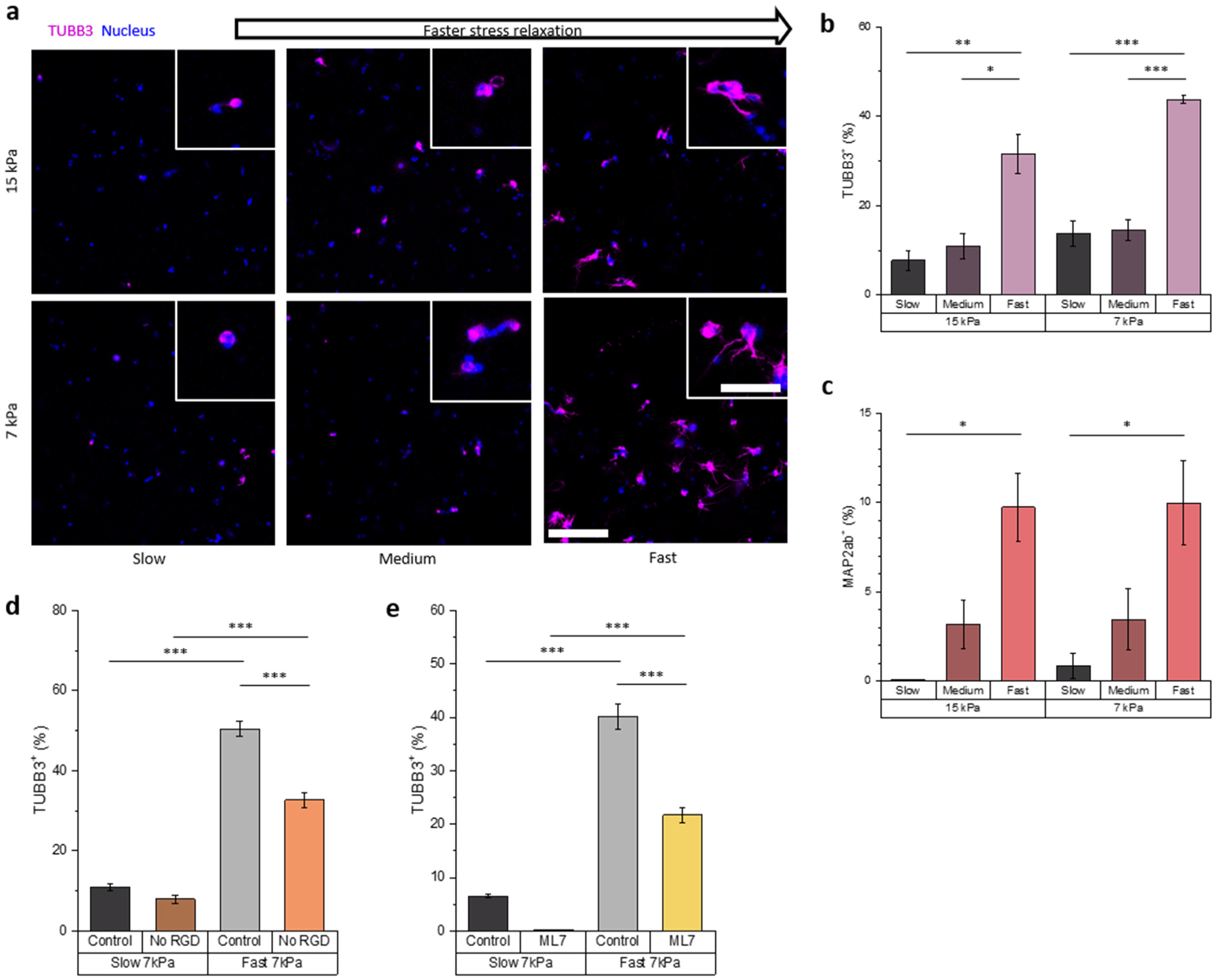
Fast matrix stress relaxation enhances neuronal differentiation of NPSCs. (a) Representative immunocytochemistry images showing TUBB3 expression and cell morphology in hydrogels with different stiffness and stress relaxation properties. Magenta, TUBB3; blue, nuclei. (b) Quantification of TUBB3-positive cell population (N = 3). (c) Quantification of MAP2ab-positive cell population (N = 3). (d) Quantification of TUBB3-positive cell population in slow- and fast-relaxing hydrogels with or without RGD ligands (N = 3). (e) Quantification of TUBB3-positive cell population in slow- and fast-relaxing hydrogels with or without ML7 treatment (N = 3). *p < 0.05, **p < 0.01, ***p < 0.001, by ANOVA with Tukey post-hoc test. Data are shown as means ± s.e.m. Scale bars: 150 µm; 50 µm in insets.

We next examined whether integrin-mediated adhesion and actomyosin contractility contribute to stress relaxation-dependent neuronal differentiation. To assess the role of integrin binding, NPSCs were cultured in slow- and fast-relaxing hydrogels with or without RGD ligands during neuronal differentiation. In fast-relaxing hydrogels, removal of RGD significantly reduced the percentage of TUBB3-positive cells, although TUBB3 expression remained above the level observed in slow-relaxing hydrogels (Fig. 2d). Similarly, inhibition of MLCK with ML7 significantly reduced neuronal differentiation in fast-relaxing hydrogels, while TUBB3 expression remained above the level observed in slow-relaxing hydrogels (Fig. 2e). These results indicate that the effect of matrix stress relaxation on NPSC neuronal differentiation is at least in part mediated by integrin binding and actomyosin contractility.,.

### 1.4. Fast matrix stress relaxation enhances astrocytic differentiation of NPSCs

To investigate how matrix stress relaxation regulates astrocytic differentiation, NPSCs were cultured in the alginate hydrogels and induced toward astrocytic differentiation using astrocyte differentiation medium for 10 days (38). Because SVZ-derived NPSCs can give rise to radial glial-like cells that express GFAP in viscoelastic matrices (Fig. 1d, e), GFAP expression alone may not be sufficient to define astrocytic differentiation in this system. We therefore used the nestin^−^GFAP^+^ population to identify astrocytes. This population expressed the astrocyte marker ALDOC and lacked the radial glial marker RC2, validating its use as an astrocytic differentiation readout in our 3D culture system (Fig. S4). NPSCs cultured in fast-relaxing hydrogels showed a significantly higher nestin^−^GFAP^+^ population and a lower nestin-positive population than those cultured in slow-relaxing hydrogels (Fig. 3a-c). This trend was observed at both stiffness levels, whereas stiffness within the tested range did not significantly affect astrocytic differentiation (Fig. 3b, c). In addition, GFAP-positive cells in faster-relaxing hydrogels showed longer processes and more primary branches, as quantified by Sholl analysis and branch-number analysis (Fig. 3h, i). These results indicate that faster matrix stress relaxation enhances astrocytic differentiation and morphological maturation under astrocyte-inducing conditions.

**Figure 3.**
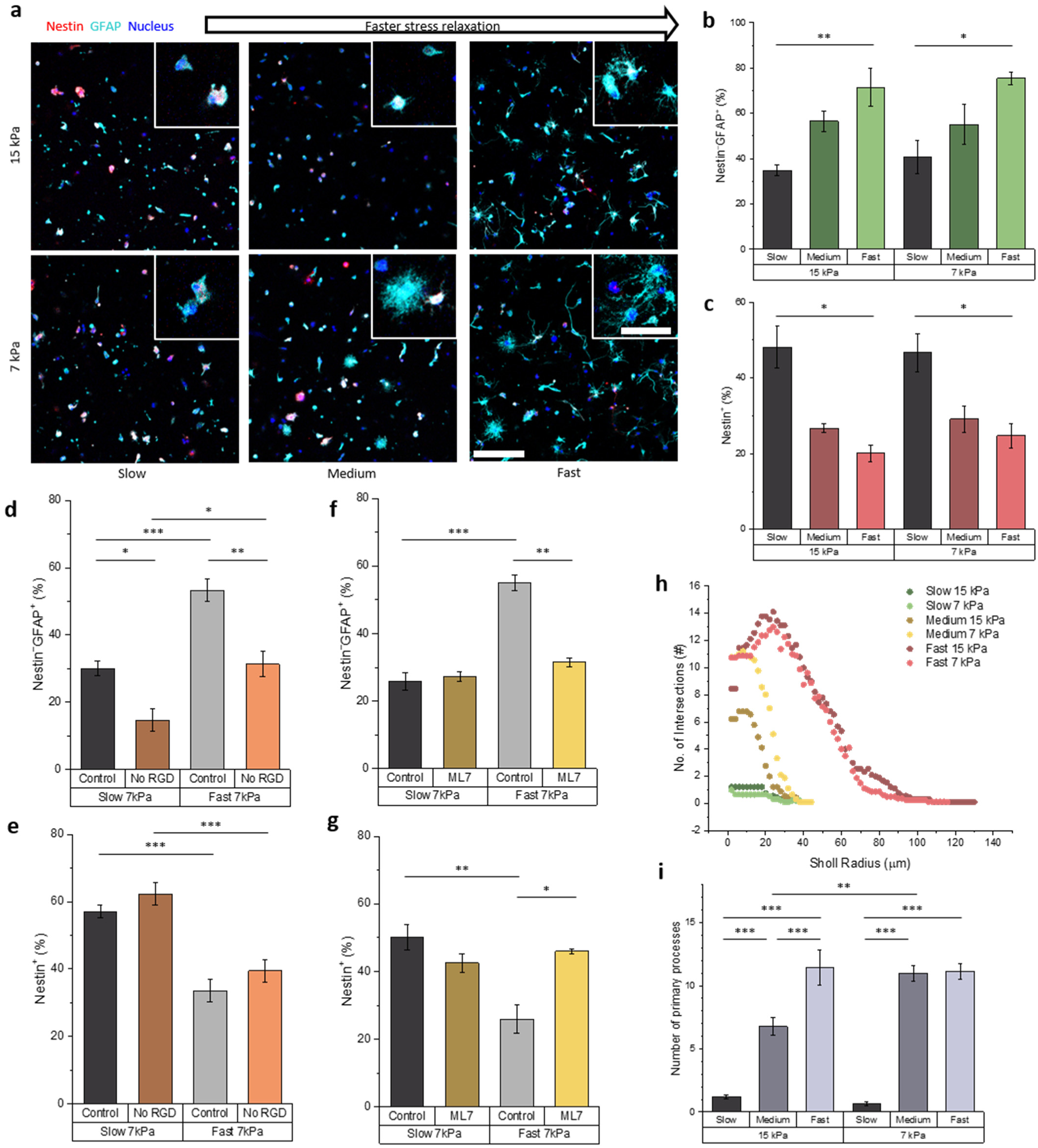
Fast matrix stress relaxation enhances astrocytic differentiation of NPSCs. (a) Representative immunocytochemistry images showing nestin and GFAP expression and cell morphology in hydrogels with different stiffness and stress relaxation properties under astrocyte-inducing conditions. Red, nestin; cyan, GFAP; blue, nuclei. (b) Quantification of nestin−GFAP^+^ cell population (N = 6). (c) Quantification of nestin-positive cell population (N = 6). (d) Quantification of nestin^−^GFAP^+^ cell population in slow- and fast-relaxing hydrogels with or without RGD ligands (N = 6). (e) Quantification of nestin-positive cell population in slow- and fast-relaxing hydrogels with or without RGD ligands (N = 6). (f) Quantification of nestin^−^GFAP^+^ cell population in slow- and fast-relaxing hydrogels with or without ML7 treatment (N = 3). (g) Quantification of nestin-positive cell population in slow- and fast-relaxing hydrogels with or without ML7 treatment (N = 3). (h) Sholl analysis of GFAP-positive cell processes (N = 9). (i) Quantification of primary branch number from GFAP-positive cell morphology analysis (N = 9). *p < 0.05, **p < 0.01, ***p < 0.001, by ANOVA with Tukey post-hoc test. Data are shown as means ± s.e.m. Scale bars: 150 µm; 50 µm in insets.

We next examined whether integrin-mediated adhesion and actomyosin contractility contribute to stress relaxation-dependent astrocytic differentiation. To assess the role of integrin binding, NPSCs were cultured in slow- and fast-relaxing hydrogels with or without RGD ligands during astrocytic differentiation. Removal of RGD significantly reduced the nestin^−^GFAP^+^ population in both slow- and fast-relaxing hydrogels, with a more pronounced reduction in fast-relaxing hydrogels (Fig. 3d). However, the nestin-positive progenitor population did not significantly change in hydrogels without RGD ligands (Fig. 3e). These results suggest that integrin-mediated adhesion has a limited effect on the loss of the progenitor marker nestin during the early stages of astrocytic differentiation, but plays a more important role at a later stage associated with acquisition of the nestin^−^GFAP^+^ astrocytic phenotype, particularly in fast-relaxing hydrogels.

To assess the role of actomyosin contractility, we inhibited MLCK using ML7. In fast-relaxing hydrogels, MLCK inhibition decreased the nestin^−^GFAP^+^ population and increased the nestin-positive population to levels comparable to those observed in slow-relaxing hydrogels (Fig. 3f, g). This indicates that actomyosin contractility is required for fast stress relaxation to enhance NPSC astrocytic differentiation. Collectively, these results suggest that integrin-mediated adhesion appears to contribute primarily to acquisition of the nestin^−^GFAP^+^ astrocytic phenotype, whereas actomyosin contractility plays a broader role in the stress relaxation-dependent differentiation response.

### 1.5. Fast matrix stress relaxation enhances oligodendrocytic differentiation of NPSCs

To investigate how matrix stress relaxation regulates oligodendrocytic differentiation, NPSCs were cultured in the alginate hydrogels and induced toward oligodendrocytic differentiation using a 7-day protocol involving sequential PDGF-AA and T3 treatment (39, 40). NPSCs cultured in fast-relaxing hydrogels showed a significantly higher O4-positive oligodendrocyte population than those cultured in slow- or medium-relaxing hydrogels (Fig. 4a, b). O4-positive cells also showed greater spreading in fast-relaxing hydrogels than those in medium- or slow-relaxing hydrogels (Fig. 4c). In contrast, stiffness within the tested range did not significantly affect either O4 expression or cell spreading (Fig. 4b, c). These results show that matrices with faster stress relaxation enhance oligodendrocytic differentiation and the spreading of NPSC-derived oligodendrocytes.

**Figure 4.**
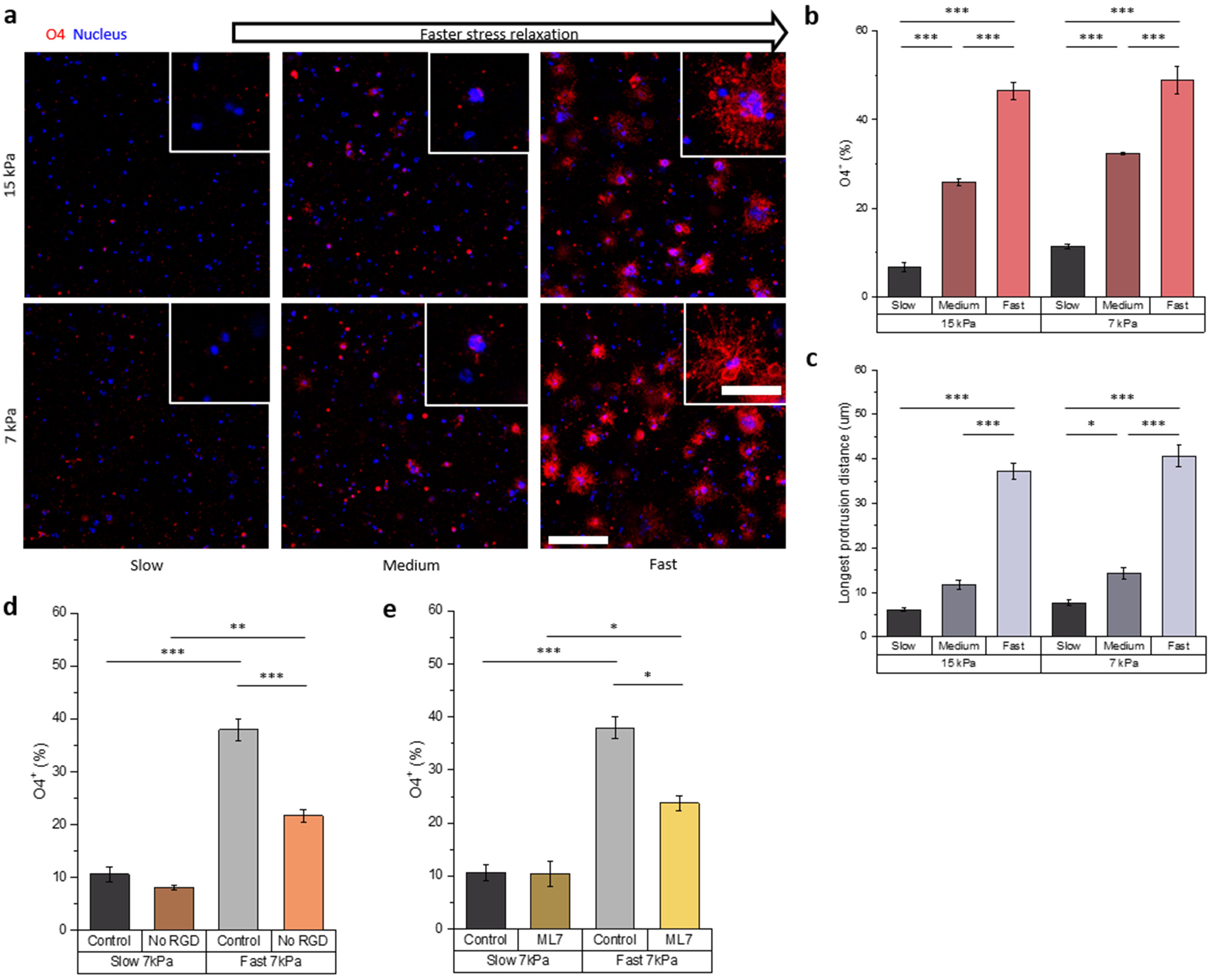
Fast matrix stress relaxation enhances oligodendrocytic differentiation outcomes. (a) Representative immunocytochemistry images showing O4 positive population and cell morphology in hydrogels with different stiffness and stress relaxation properties under oligodendrocyte-inducing conditions. Red, O4; blue, nuclei. (b) Quantification of O4-positive cell population (N = 3). (c) Quantification of the longest protrusion distance from the center of the soma or cluster (N = 35). (d) Quantification of O4-positive cell population in slow- and fast-relaxing hydrogels with or without RGD ligands (N = 3). (e) Quantification of O4-positive cell population in slow- and fast-relaxing hydrogels with or without ML7 treatment (N = 3). *p < 0.05, **p < 0.01, ***p < 0.001, by ANOVA with Tukey post-hoc test. Data are shown as means ± s.e.m. Scale bars: 150 µm; 50 µm in insets.

We next examined whether integrin-mediated adhesion and actomyosin contractility contribute to stress relaxation-dependent oligodendrocytic differentiation. To assess the role of integrin binding, NPSCs were cultured in slow- and fast-relaxing hydrogels with or without RGD ligands during oligodendrocytic differentiation. In slow-relaxing hydrogels, removal of RGD did not significantly affect the O4-positive population. In contrast, in fast-relaxing hydrogels, removal of RGD significantly reduced the O4-positive population, although the O4-positive population remained above the level observed in slow-relaxing hydrogels (Fig. 4d). These results indicate that integrin-mediated adhesion contributes to the enhancement of oligodendrocytic differentiation induced by fast matrix stress relaxation, but does not fully account for this effect. To assess the role of actomyosin contractility, we inhibited MLCK using ML7. MLCK inhibition significantly reduced the O4-positive population in fast-relaxing hydrogels, while having no significant effect in slow-relaxing hydrogels (Fig. 4e). These findings indicate that actomyosin contractility also contributes to the stress-relaxation-dependent enhancement of oligodendrocytic differentiation.

### 1.6. Fast matrix stress relaxation preferentially promotes neuronal differentiation under mixed neuronal/astrocytic differentiation conditions

To investigate whether matrix stress relaxation biases NPSC differentiation under mixed neuronal and astrocytic cues, NPSCs were cultured in the alginate hydrogels and treated with medium containing retinoic acid and FBS to provide cues for both neuronal and astrocytic differentiation (41). Under these mixed differentiation conditions, NPSCs cultured in fast-relaxing hydrogels showed a higher proportion of TUBB3-positive cells than those cultured in slow- or medium-relaxing hydrogels at both stiffness levels (Fig. 5a, b). In contrast, the GFAP-positive astroglial population showed an opposite trend, with lower levels in faster-relaxing hydrogels (Fig. 5c). Nestin-positive population showed a decreasing trend with faster stress relaxation, although this change was not statistically significant (Fig. 5d). These results indicate that, under mixed neuronal/astrocytic differentiation conditions, fast matrix stress relaxation preferentially promotes neuronal differentiation relative to astrocytic differentiation.

**Figure 5.**
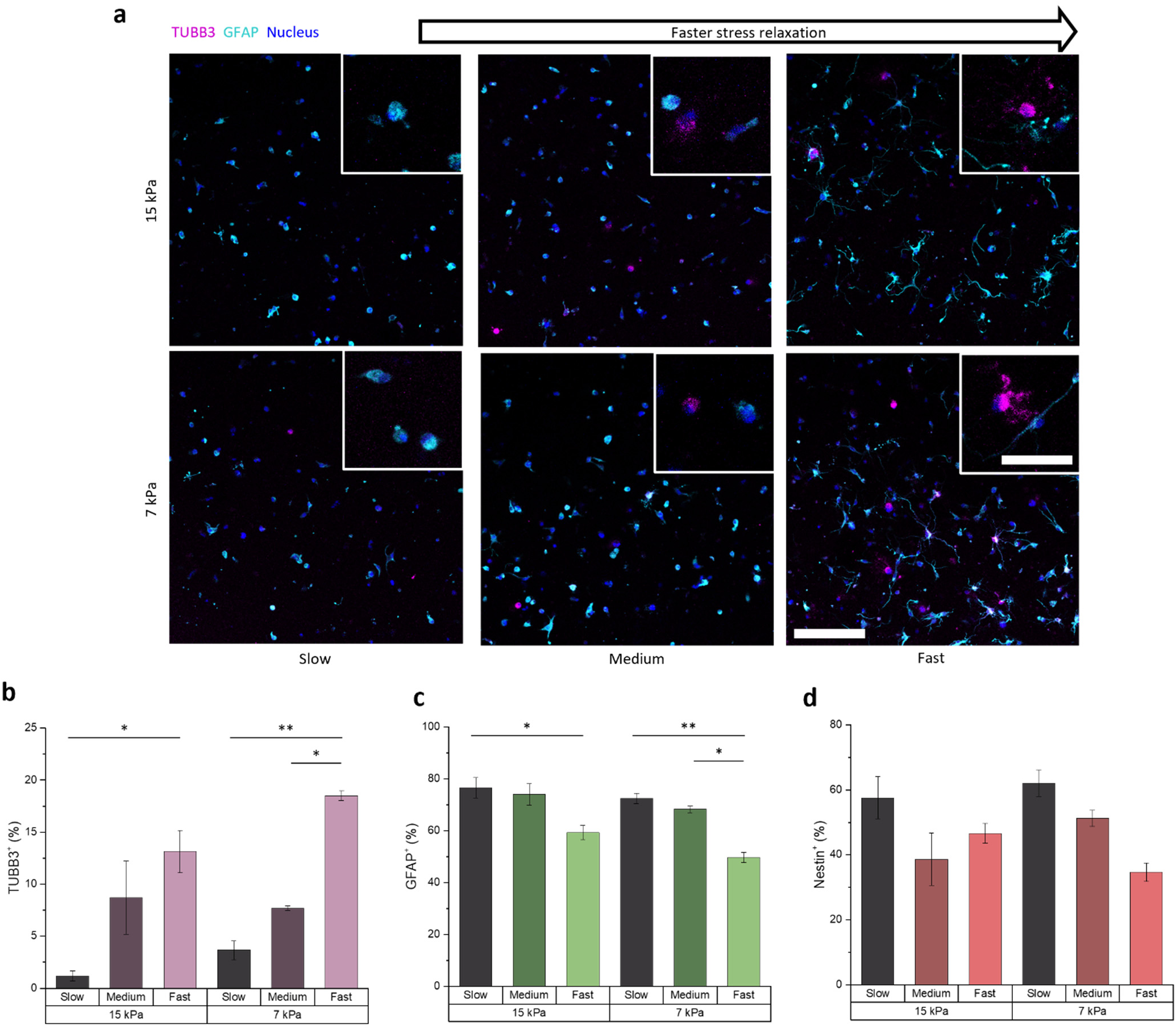
Fast matrix stress relaxation biases mixed neuronal/astrocytic differentiation toward neuronal outcomes. (a) Representative immunocytochemistry images showing TUBB3 and GFAP expression and cell morphology in hydrogels with different stiffness and stress relaxation properties under mixed neuronal/astrocytic differentiation conditions. Magenta, TUBB3; cyan, GFAP; blue, nuclei. (b) Quantification of TUBB3-positive cell population (N = 3). (c) Quantification of GFAP-positive astroglial population (N = 3). (d) Quantification of nestin-positive cell population (N = 3). *p < 0.05, **p < 0.01, by ANOVA with Tukey post-hoc test. Data are shown as means ± s.e.m. Scale bars: 150 µm; 50 µm in insets.

### 1.7. Matrix stress relaxation regulates neuronal/astrocytic fate bias through integrin-mediated adhesion, actomyosin contractility, and actin polymerization

We next examined whether integrin-mediated adhesion and cytoskeletal mechanotransduction pathways contribute to the stress-relaxation-dependent fate bias observed under mixed neuronal/astrocytic differentiation conditions. To assess the role of integrin binding, NPSCs were cultured in slow- and fast-relaxing hydrogels with or without RGD ligands during mixed differentiation. In slow-relaxing hydrogels, removal of RGD did not significantly affect either the TUBB3-positive neuronal population or the GFAP-positive astroglial population (Fig. 6b, c). In contrast, in fast-relaxing hydrogels, removal of RGD significantly reduced the TUBB3-positive population and increased the GFAP-positive astroglial population (Fig. 6b, c). In fast-relaxing 15 kPa hydrogels, removal of RGD reduced the TUBB3-positive proportion to a level comparable to that observed in slow-relaxing hydrogels, whereas in fast-relaxing 7 kPa hydrogels, the TUBB3- positive proportion remained above the level observed in slow-relaxing hydrogels (Fig. 6b). These results indicate that integrin-mediated adhesion contributes to the stress-relaxation-dependent neuronal bias under mixed differentiation conditions, with a stronger effect in the 15 kPa condition.

**Figure 6.**
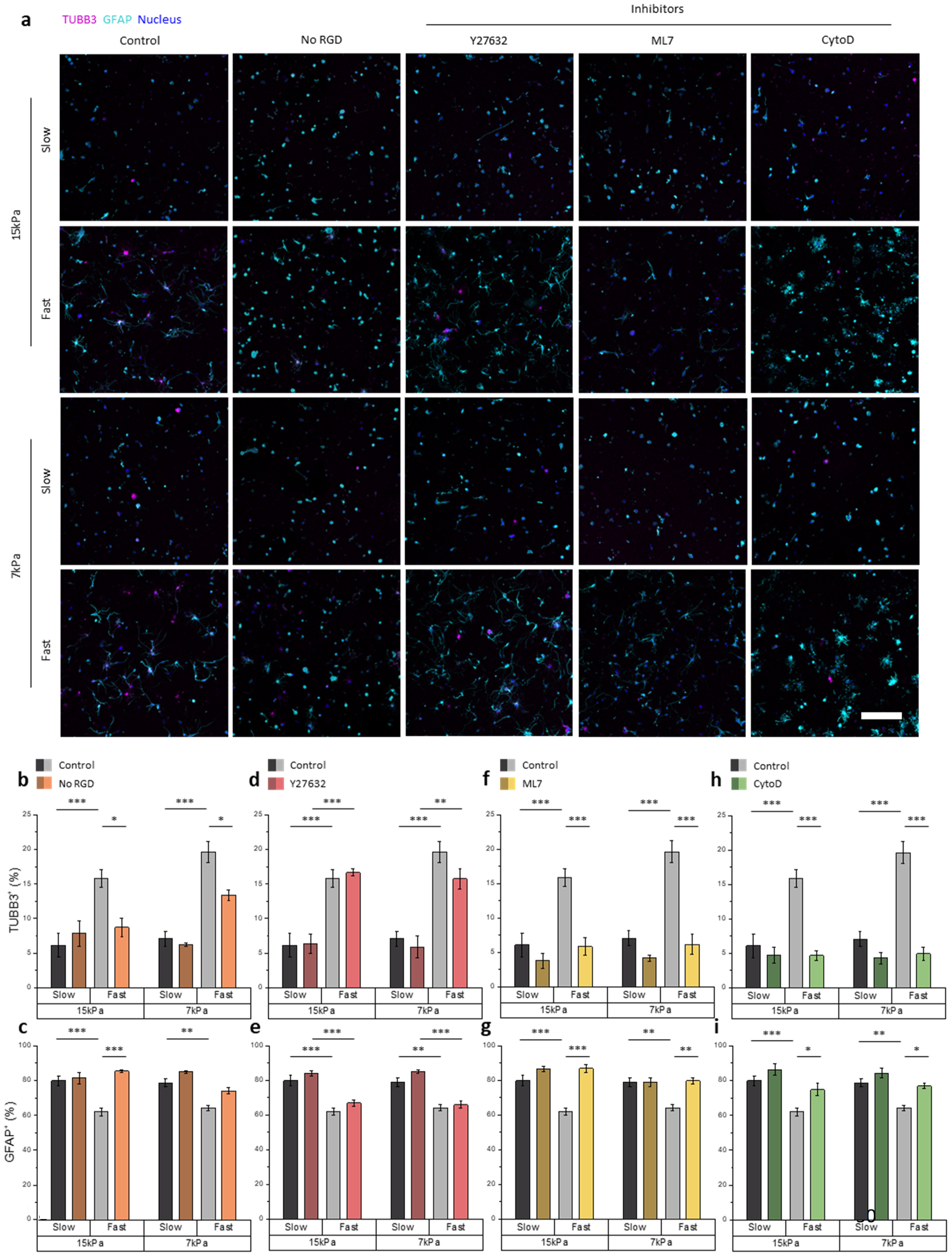
Matrix stress relaxation regulates neuronal/astrocytic fate bias through integrin-mediated adhesion, actomyosin contractility, and actin polymerization. (a) Representative immunocytochemistry images showing TUBB3 and GFAP expression and cell morphology in hydrogels with different stiffness and stress relaxation properties under mixed neuronal/astrocytic differentiation conditions, with or without RGD ligands or cytoskeletal inhibitors. Magenta, TUBB3; cyan, GFAP; blue, nuclei. (b, c) Quantification of TUBB3-positive neuronal population and GFAP-positive astroglial population in slow- and fast-relaxing hydrogels with or without RGD ligands (N = 9). (d, e) Quantification of TUBB3-positive neuronal population and GFAP-positive astroglial population in slow- and fast-relaxing hydrogels with or without Y-27632 treatment (N = 7). (f, g) Quantification of TUBB3-positive neuronal population and GFAP-positive astroglial population in slow- and fast-relaxing hydrogels with or without ML7 treatment (N = 8). (h, i) Quantification of TUBB3-positive neuronal population and GFAP-positive astroglial population in slow- and fast-relaxing hydrogels with or without cytochalasin-D (CytoD) treatment (N = 7). *p < 0.05, **p < 0.01, ***p < 0.001, by ANOVA with Tukey post-hoc test. Data are shown as means ± s.e.m. Scale bar: 150 µm.

We next tested whether actomyosin contractility and actin polymerization contribute to this response. Inhibition of ROCK using Y-27632 did not significantly affect the TUBB3-positive or GFAP-positive astroglial populations in any of the tested mechanical conditions (Fig. 6d, e). In contrast, inhibition of MLCK using ML7 significantly reduced TUBB3-positive proportion and increased the GFAP-positive astroglial population in fast-relaxing hydrogels, while having no significant effect in slow-relaxing hydrogels (Fig. 6f, g). Similarly, inhibition of actin polymerization using cytochalasin-D reduced TUBB3-positive proportion and increased the GFAP-positive astroglial population in fast-relaxing hydrogels, but did not significantly affect mixed differentiation outcomes in slow-relaxing hydrogels (Fig. 6h, i). These inhibitor effects were also reflected in cell morphology, with ML7 and cytochalasin-D visibly reducing the spreading morphology observed in fast-relaxing hydrogels, whereas ROCK inhibition had limited morphological effect (Fig. 6a).

These results show that the stress relaxation-dependent neuronal bias under mixed neuronal/astrocytic differentiation conditions is regulated by integrin-mediated adhesion, MLCK-dependent actomyosin contractility, and actin polymerization, but is not significantly affected by ROCK inhibition. This indicates that downstream cytoskeletal regulation, rather than ROCK activity alone, contributes to the effect of fast matrix stress relaxation on NPSC fate bias under mixed differentiation cues.

### 1.8. The effect of matrix stress relaxation on neuronal/astrocytic fate bias is also mediated by Piezo1 activity

We next examined whether the mechanosensitive ion channel Piezo1 mediates matrix stress relaxation-dependent regulation of NPSC fate under mixed neuronal/astrocytic differentiation conditions. Piezo1 expression was assessed in NPSCs 3D-cultured in stress-relaxing hydrogels using immunofluorescence staining followed by flow cytometry. Under 7 kPa conditions, NPSCs had significantly higher Piezo1 expression in slow-relaxing gels than in fast-relaxing gels. The difference at 15 kPa showed similar trend but did not reach statistical significance (Fig. S5a).

To investigate whether Piezo1 activity affects mixed differentiation outcomes, we treated NPSCs with the Piezo1 inhibitor GsMTx4 or the Piezo1 agonist Yoda1 during mixed neuronal/astrocytic differentiation in 7 kPa hydrogels (Fig. 7a). In fast-relaxing hydrogels, GsMTx4 significantly reduced the proportion of TUBB3-positive cells, whereas Yoda1 significantly increased this proportion relative to their respective controls (Fig. 7b,d). Neither treatment significantly altered the proportion of TUBB3-positive cells in slow-relaxing hydrogels (Fig. 7b,d). The proportion of GFAP-positive astroglial cells was not significantly altered by either treatment under either relaxation condition (Fig. 7c,e). Corresponding experiments in 15 kPa hydrogels showed changes in TUBB3-positive cell proportions in the same respective directions in fast-relaxing matrices, although these treatment effects did not reach statistical significance (Fig. S5b-f). Together, these findings support a role for Piezo1 activity in regulating neuronal differentiation in fast-relaxing hydrogels under mixed neuronal/astrocytic differentiation conditions.

**Figure 7.**
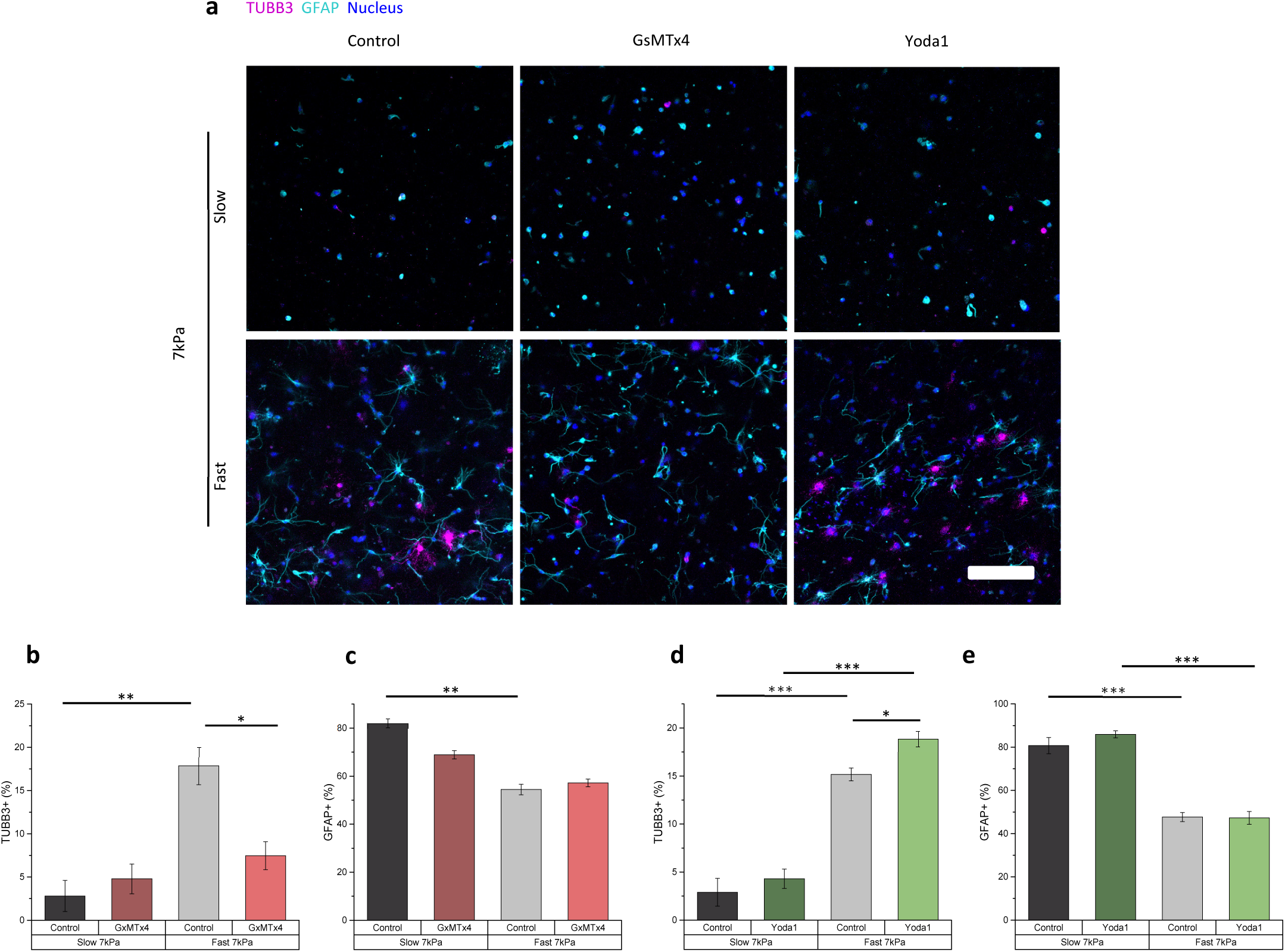
Piezo1 inhibition and activation influence the effect of matrix stress relaxation on the neuronal differentiation under mixed neuronal/astrocytic differentiation conditions. (a) Representative immunocytochemistry images showing TUBB3 and GFAP expression and cell morphology in hydrogels with different stiffness and stress relaxation properties under mixed neuronal/astrocytic differentiation conditions, with or without the Piezo1 inhibitor GsMTx4 or the Piezo1 agonist Yoda1. Magenta, TUBB3; cyan, GFAP; blue, nuclei. (b, c) Quantification of TUBB3-positive cell population and GFAP-positive astroglial population with or without GsMTx4 treatment (N = 3). (d, e) Quantification of TUBB3-positive cell population and GFAP-positive astroglial population with or without Yoda1 treatment (N = 3). *p < 0.05, **p < 0.01, ***p < 0.001, by ANOVA with Tukey post-hoc test. Data are shown as means ± s.e.m. Scale bar: 150 µm.

## 2. Discussion

In this study, we show that 3D matrix stress relaxation regulates the stemness and multilineage fate potential of primary NPSCs. Using alginate hydrogels with independently tunable stiffness and stress relaxation, we found that faster matrix stress relaxation enhanced radial glial-like marker expression, NPSC stemness maintenance, and differentiation toward neuronal, astrocytic, and oligodendrocytic lineages under corresponding biochemical induction conditions. Under mixed neuronal/astrocytic differentiation conditions, fast-relaxing matrices preferentially promoted neuronal differentiation relative to the GFAP-positive astroglial population. These effects were observed consistently across two stiffness levels. Mechanistically, NPSC responses to matrix stress relaxation involved integrin-mediated adhesion, MLCK-dependent actomyosin contractility, actin polymerization, and Piezo1 activity, with distinct contributions depending on the differentiation context. Together, these findings establish matrix stress relaxation as an important 3D mechanical cue that regulates primary NPSC maintenance, lineage specification, and fate bias through context-dependent mechanotransduction (Fig. 8).

**Figure 8.**
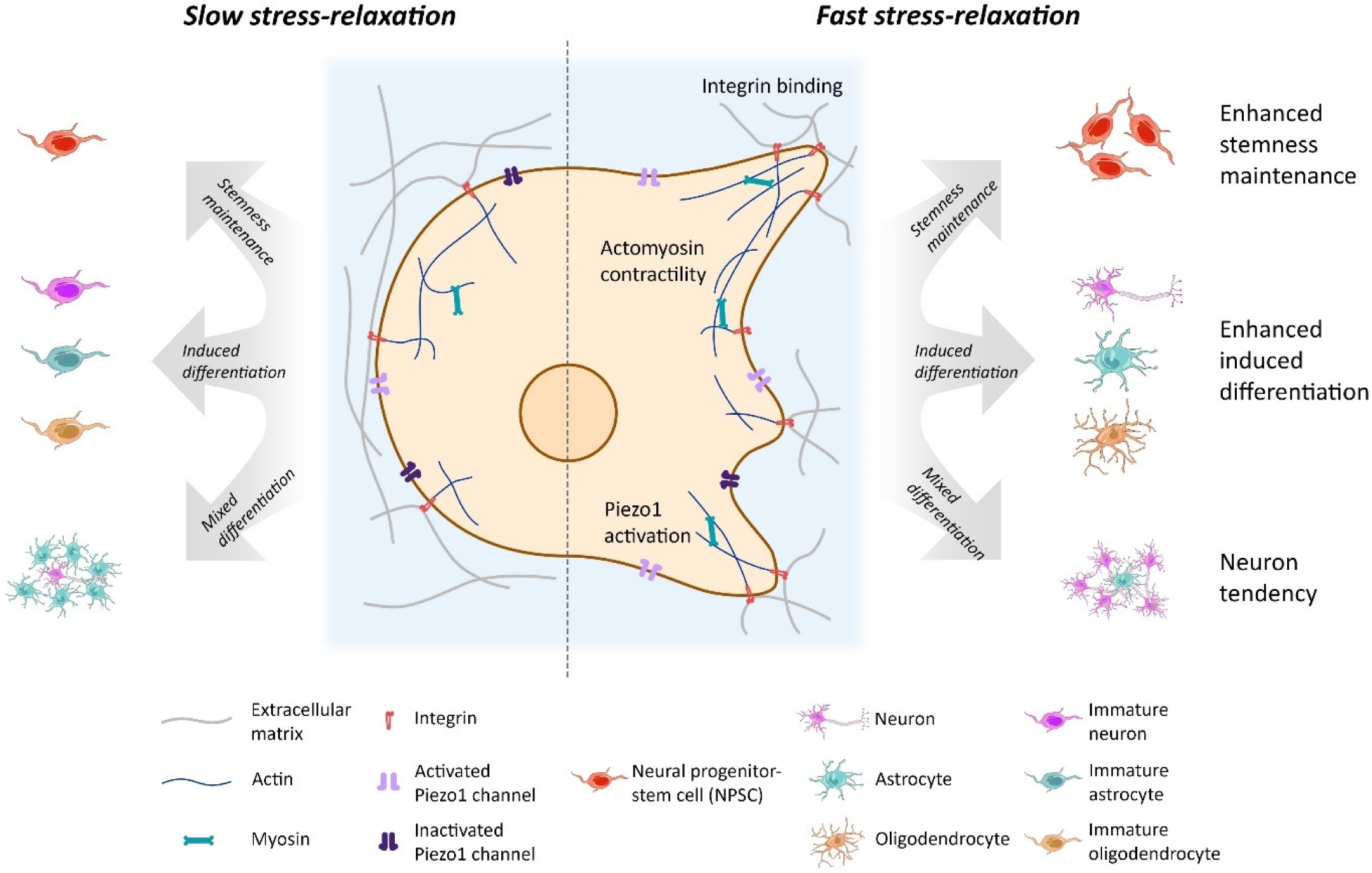
Schematic illustration of NPSC stemness maintenance and differentiation regulated by matrix stress relaxation. Fast-relaxing matrices support NPSC stemness maintenance and enhance induced differentiation under lineage-specific biochemical conditions. Under mixed neuronal/astrocytic differentiation conditions, fast-relaxing matrices preferentially promote neuronal differentiation relative to astrocytic differentiation. These responses involve matrix-cell mechanotransduction pathways including integrin-mediated adhesion, actomyosin contractility, actin polymerization, and Piezo1 activity.

One notable finding is that 3D culture in viscoelastic hydrogels supported the re-emergence of radial glial-like progenitor markers and enhanced stemness maintenance in SVZ-derived NPSCs. After 2D expansion, the cells retained nestin expression but showed little or no GFAP and RC2 expression, consistent with previous reports that radial glial marker expression can change with developmental stage and in vitro culture conditions (33–35). Subsequent culture in 3D viscoelastic hydrogels restored GFAP and RC2 expression, especially in fast-relaxing matrices (Fig. 1d, e). Because radial glial-like progenitors in the SVZ are associated with nestin, GFAP, and RC2 expression, the appearance of nestin^+^GFAP^+^RC2^+^ and nestin^+^GFAP^−^RC2^+^ populations suggests that the 3D stress-relaxing environment helps maintain or recover progenitor marker states related to the endogenous SVZ niche (24, 31, 32). Consistent with this interpretation, faster-relaxing hydrogels also promoted greater cell spreading and aggregation (Fig. 1f), increased cell number (Fig. 1g), and maintained a higher nestin-positive population than slow-relaxing hydrogels (Fig. 1h). These effects were observed at both stiffness levels. Because the quantity of calcium sulfate crosslinker was varied to tune hydrogel mechanics, we further examined whether the stemness-maintenance response could instead be explained by initial Ca²⁺ crosslinker concentration. When nestin-positive proportion was replotted against initial Ca²⁺ crosslinker concentration, the response did not show an obvious correlation with Ca²⁺ concentration, but instead aligned more closely with stress-relaxation condition (Fig. S2).

Together with the RGD-removal and MLCK-inhibition results, which indicate that integrin-mediated adhesion and actomyosin contractility are required for fast stress relaxation to enhance nestin expression (Fig. 1i, j), these findings support the interpretation that faster-relaxing 3D matrices enhance stemness maintenance through mechanical matrix engagement rather than through stiffness or initial Ca²⁺ crosslinker concentration alone. The influence of stiffness also differs from previous 2D studies in which softer substrates better maintain NPSC stemness or promote neuronal differentiation (42), emphasizing that mechanical regulation of NPSCs can depend strongly on dimensionality and matrix architecture. The stemness-supporting effect of fast stress relaxation is also consistent with prior work showing that 3D matrix remodeling via biochemical degradation supports NPSC stemness maintenance and differentiation capacity (17, 19), as well as with neurosphere studies suggesting that 3D organization and viscoelastic properties may contribute to NPSC maintenance in addition to biochemical cell-cell interactions (43–45). Together, these findings suggest that fast-relaxing 3D matrices provide a permissive mechanical environment that supports SVZ-derived NPSC progenitor features, cell expansion/survival, and nestin-positive stemness maintenance.

Fast matrix stress relaxation promoted neuronal outcomes under both defined neurogenic and mixed neuronal/astrocytic differentiation conditions. Under the neuronal differentiation protocol, fast-relaxing hydrogels increased the TUBB3-positive population and enhanced MAP2ab expression, indicating increased neuronal differentiation and maturation under neurogenic cues (Fig. 2b, c; Fig. S3). Under mixed neuronal/astrocytic differentiation conditions, fast-relaxing matrices again increased the TUBB3-positive population while reducing the GFAP-positive astroglial population (Fig. 5b, c). This indicates faster stress relaxation can enhance neuronal differentiation across distinct biochemical environments. This finding is consistent with recent studies showing that matrix viscoelasticity can promote neurogenic or neuronal maturation outcomes. In adult hippocampal NSCs, stress-relaxing 3D HA matrices increased β-tubulin III-positive neurogenesis and reduced GFAP-positive astrogenesis under mixed differentiation conditions, whereas substrate stress relaxation produced the opposite trend in 2D culture (12, 13). Tunable hydrogel viscoelasticity has also been shown to promote human neural maturation and neurite extension through actin-polymerization-dependent mechanisms (21). However, in engineered ELP-PEG hydrogels, faster stress relaxation promoted both neuronal and astroglial differentiation capacity (23), highlighting a parallel enhancement rather than the neuronal/astroglial competition mechanism in our result. This suggests the dependence of chemical environment, which is further supported by the glial lineage-specific differentiation studies discussed below.

Under astrocyte-inducing conditions, fast-relaxing matrices enhanced astrocytic differentiation, in contrast to the reduced astroglial population observed under mixed neuronal/astrocytic differentiation conditions. Fast-relaxing hydrogels increased the nestin^−^GFAP^+^ population, reduced the nestin-positive progenitor population, and promoted longer and more branched GFAP-positive cell morphologies (Fig. 3b, c, h, i). Because SVZ-derived NPSCs can re-express GFAP as part of a radial glial-like progenitor marker profile in 3D stress-relaxing hydrogels, the nestin^−^GFAP^+^ population, supported by ALDOC expression and lack of RC2 expression, provides a more specific readout of astrocytic differentiation in this system (Fig. S4). Similarly, under oligodendrocyte-inducing conditions, fast stress relaxation enhanced O4-positive oligodendrocytic differentiation outcomes, as shown by an increased O4-positive population and greater O4-positive cell spreading in fast-relaxing hydrogels (Fig. 4b, c). Previous studies have shown that oligodendrocyte lineage cells respond to matrix stiffness and mechanical strain (46–50). Collectively, these results indicate that fast matrix stress relaxation does not impose a single fixed lineage outcome, but instead enhances NPSC responsiveness to lineage-specific biochemical cues, including astrocytic and oligodendrocytic induction.

The mechanotransduction data indicate that integrin-mediated adhesion and MLCK-dependent actomyosin contractility are important but not universally equivalent mediators of stress-relaxation-dependent NPSC responses. For stemness maintenance, removal of RGD ligands or inhibition of MLCK reduced nestin expression in fast-relaxing hydrogels to levels comparable to those in slow-relaxing hydrogels, suggesting that integrin engagement and actomyosin contractility are required for fast stress relaxation to enhance stemness maintenance (Fig. 1i, j). In contrast, during neuronal and oligodendrocytic differentiation, RGD removal and MLCK inhibition reduced the fast-relaxation effect but did not fully eliminate it, indicating that these pathways contribute to, but do not fully account for, stress-relaxation-dependent differentiation under these lineage-inducing conditions (Fig. 2d, e and Fig. 4d, e). During astrocytic differentiation, integrin-mediated adhesion appeared to contribute more strongly to acquisition of the nestin^−^GFAP^+^ astrocytic phenotype than to early loss of nestin expression, whereas MLCK inhibition more broadly suppressed the fast-relaxation-dependent astrocytic differentiation response (Fig. 3d-g). These differences suggest that NPSCs do not use a single mechanotransduction pathway to respond to matrix stress relaxation cues. This interpretation is consistent with emerging studies showing that viscoelastic matrices can regulate neural progenitor behavior through multiple mechanisms, including actin polymerization, spectrin, N-cadherin-like interactions, and β-catenin-related signaling (12, 13, 21–23).

The mixed neuronal/astrocytic differentiation condition revealed a broader mechanotransduction network regulating stress-relaxation-dependent fate bias. Under these conditions, RGD removal, MLCK inhibition, and actin polymerization inhibition reduced the TUBB3-positive population and increased the GFAP-positive astroglial population in fast-relaxing hydrogels, indicating that integrin-mediated adhesion, actomyosin contractility, and actin polymerization contribute to the neuronal bias induced by fast stress relaxation (Fig. 6b, c, f-i). In contrast, ROCK inhibition did not significantly alter the TUBB3-positive or GFAP-positive astroglial populations, despite the strong effects of MLCK and actin polymerization inhibition (Fig. 6d, e). Piezo1 activity also contributed to neuronal differentiation under mixed differentiation conditions. Piezo1 expression varied across stress-relaxing matrix conditions (Fig. S5), and pharmacological modulation of Piezo1 selectively altered the TUBB3-positive population in fast-relaxing 7 kPa hydrogels: GsMTx4 reduced TUBB3-positive proportion, whereas Yoda1 increased TUBB3-positive proportion(Fig. 7b, d). In contrast, the GFAP-positive astroglial population was not significantly altered by either treatment (Fig. 7c, e). These findings suggest that Piezo1 acts as a context-specific mechanotransduction element regulating neuronal differentiation. Because Piezo1 has been implicated in coupling extracellular mechanical cues to intracellular signaling and neural stem/progenitor cell fate regulation (26–28, 51, 52), these results support a model in which fast matrix stress relaxation regulates neuronal differentiation through coordinated mechanosensitive ion-channel activity and cytoskeletal remodeling. Future studies directly measuring Piezo1 activation dynamics, downstream signaling, and cytoskeletal force transmission will be needed to define how these pathways interact during stress-relaxation-dependent NPSC fate regulation.

Several considerations frame the interpretation and future extension of this work. The use of primary SVZ-derived NPSCs provides a more physiologically relevant extension of studies using immortalized neural progenitor lines or pluripotent stem cell-derived neural models, because these cells are isolated from an endogenous neurogenic niche and retain multilineage differentiation potential. However, NPSC responses to matrix stress relaxation may vary with developmental stage, species, and progenitor source, making it important for future studies to compare primary postnatal, adult, human, and pluripotent stem cell-derived neural progenitor populations.

Overall, this study demonstrates that 3D matrix stress relaxation regulates the stemness and multilineage fate potential of primary NPSCs. Using alginate hydrogels with independently tunable stiffness and stress relaxation, we found that faster matrix stress relaxation supports radial glial-like marker expression, enhances nestin-positive stemness maintenance, and promotes neuronal, astrocytic, and oligodendrocytic differentiation of NPSCs under the corresponding biochemical induction conditions. Under mixed neuronal/astrocytic differentiation conditions, fast-relaxing matrices biased NPSC fate toward neuronal differentiation relative to the GFAP-positive astroglial population, indicating that the effect of stress relaxation depends on the biochemical differentiation environment. These responses were associated with multiple mechanotransduction pathways, including integrin-mediated adhesion, MLCK-dependent actomyosin contractility, actin polymerization, and Piezo1 activity, with distinct contributions across stemness maintenance, lineage-specific differentiation, and mixed-fate regulation.

## 3. Materials and Methods

### 3.1. Alginate preparation

Sodium alginate with high guluronic acid block content and high molecular weight (high-MW, 153.8 kDa, LF20/40) was purchased from FMC Biopolymer and further irradiated by a cobalt-60 source at 3 or 8 MRad to produce mid-MW (43.7 kDa) or low-MW (25.0 kDa) alginate (53). RGD-alginate was prepared by coupling oligopeptide GGGGRGDSP (Peptide 2.0 Inc.) with carbodiimide chemistry (29). A final RGD density of 1.5 mM was achieved in a 2% wt/vol alginate gel. Alginate was dialyzed against deionized water for 3 days using a dialysis membrane (3500 MWCO, Spectrum Chemical), purified by activated charcoal, sterile filtered, lyophilized, stored at −20 °C, and reconstituted in DMEM/F12 (Thermo Fisher Scientific) before use.

### 3.2. Mechanical characterization

Gel disks (15 mm in diameter, 2 mm in thickness) of alginate with 2% wt/vol alginate and different calcium concentrations were prepared using our previously published method (29, 54).

Unconfined uniaxial compression tests were performed on the disks at a 2 mm/min loading rate until 15% strain, followed by stress relaxation tests in which the constant strain was held. Initial elastic modulus was measured from the slope of the stress-strain curve within the 5–10% strain range, while relaxation half-time was calculated as the time required for the stress to relax from peak stress to half of the peak stress during the stress relaxation test.

### 3.3. Primary NPSC isolation and expansion

All animal experiments were performed following protocols approved by the Johns Hopkins University Animal Care and Use Committee.

The isolation and expansion of rat neural progenitor-stem cells (NPSCs) used a previously described protocol (30). Briefly, postnatal day 7 Sprague-Dawley rat (Taconic Biosciences) brains were quickly dissected into ice-cold HBSS with 2% Pen/Strep, where they were further dissected to obtain the SVZ tissues. The SVZ tissues were transferred into 37 °C 0.05% Trypsin/EDTA

(Thermo Fisher Scientific) and incubated for 10 min, after which trypsin inhibitor (Roche) and DNase I (Worthington Biochem) were added to neutralize enzymatic activity. Tissue suspension was then centrifuged at 300 xg for 5 min and the supernatant was discarded. The centrifuged SVZ tissues were resuspended in NPSC maintenance medium consisting of NeuroCult™ Proliferation Kit (Stemcell Technologies), 2 μg/mL heparin (Sigma, H3149), 20 ng/mL EGF (PeproTech), 10 ng/mL bFGF (PeproTech), and 50 U/mL Pen/Strep, and triturated approximately 10 times to obtain a cell suspension. The cell suspension was passed through a 70 μm cell strainer (BD Falcon) and the isolated cells were cultured on Matrigel-coated (Corning) 6-well plates until 90% confluent and ready for passaging. Half of the medium was changed every 2 days. Proliferative NPSCs were passaged up to passage 2 before use in subsequent experiments.

### 3.4. NPSC encapsulation in alginate

NPSCs were detached from tissue culture plates using Accutase (Thermo Fisher Scientific) for 6 min at 37 °C, resuspended in serum-free DMEM/F12 at 37.5 million cells/mL, and mixed with RGD-alginate reconstituted in DMEM/F12. The cell-alginate suspension was rapidly mixed with calcium sulfate at different concentrations in DMEM/F12, leading to a final cell density of 5 million cells/mL. The NPSC-encapsulated alginate gels were formed between two glass plates spaced by 1 mm for 30 min at room temperature. After that, the alginate gels were punched into disks (6 mm in diameter, 1 mm in thickness) and cultured in the corresponding media. Monolayer culture of NPSCs on Matrigel-coated petri dishes was performed in parallel.

### 3.5. NPSC culture and differentiation

NPSC stemness maintenance was performed by culturing NPSCs in NPSC maintenance medium for 7 days. Half of the medium was changed every 2 days.

Neuronal differentiation was performed based on a previously described protocol (37). NPSCs were cultured in DMEM/F12 containing 1% N2 and 0.5 μM retinoic acid (Sigma) for 3 days, followed by DMEM/F12 containing 1% N2 (Thermo Fisher Scientific), 0.5% FBS (Hyclone), and 20 ng/mL rhBDNF (PeproTech) for 12 days. The medium was changed every 2 days.

Astrocytic differentiation was performed based on a previously described protocol (38). NPSCs were cultured in DMEM/F12 containing 2% B27 (Thermo Fisher Scientific), 1% N2 (Thermo Fisher Scientific), and 5% FBS (Hyclone) for 10 days. The medium was changed every 2 days.

Oligodendrocytic differentiation was performed using a previously described protocol (39, 40). NPSCs were cultured in DMEM/F12 containing 1% B27, 1% N2, 20 ng/mL bFGF, 20 ng/mL PDGF-AA (PeproTech), and 0.4 μM SAG (Enzo) for 4 days, followed by complete medium change to DMEM/F12 containing 1% B27, 1% N2, 30 ng/mL T3 (Sigma), 0.4 μM SAG, 100 ng/mL noggin (R&D Systems), 10 μM cAMP (Sigma), 100 ng/mL IGF1 (PeproTech), and 10 ng/mL NT3 (PeproTech) for 3 days. The medium was changed every 3 days.

Mixed neuronal/astrocytic differentiation was performed using a previously described protocol (41, 55). NPSCs were cultured in DMEM/F12 containing 1% N2, 1 μM retinoic acid, and 1% FBS for 7 days. The medium was changed every 2 days.

Viability tests were performed before and after differentiation by staining samples with Hoechst (1:200) and propidium iodide (1:500) in Ca-HEPES buffer containing 20 mM HEPES (Alfa Aesar), 1.5 mM CaCl_2_, and 138 mM NaCl for 20 min at 37 °C.

### 3.6. Immunocytochemistry

Immunocytochemistry was performed to evaluate stemness and differentiation outcomes. For staining of cells cultured either in alginate or on petri dishes, samples were first washed once with Ca-HEPES buffer and fixed with 4% paraformaldehyde in Ca-HEPES buffer for 30 min at room temperature. The samples were then washed three times with Ca-HEPES buffer, permeabilized with 0.2% Triton X-100 (Thermo Fisher Scientific) in Ca-HEPES buffer for 20 min, and blocked with 2.5% BSA (Sigma) in Ca-HEPES buffer for 2 h at room temperature. Primary antibody staining was performed overnight at 4 °C. After three washes, secondary antibody staining was performed together with Hoechst (1:200) for 2 h at room temperature. When donkey anti-goat secondary antibody was used with other goat-source secondary antibodies on the same samples, donkey anti-goat secondary antibody was applied for 2 h first and washed three times, followed by staining with other secondary antibodies for 2 h. All antibodies were diluted in Ca-HEPES buffer for staining. Cells cultured on petri dishes were imaged using an EVOS M5000 microscope, while cells cultured in alginate were imaged as 50 μm z-stacks using an LSM800 confocal microscope.

The following primary antibodies were used for immunostaining: rabbit anti-rat ALDOC (1:200, Proteintech, 14884-1-AP), mouse anti-rat Aqp4 (1:400, Millipore Sigma, MABN2527), rabbit anti-rat GFAP (1:1000, Dako, Z0334), rat anti-rat GFAP (1:500, Invitrogen, 13-0300), mouse anti-rat MAP2ab (1:200, Invitrogen, MA5-12823), goat anti-rat nestin (1:80, R&D Systems, AF2736), mouse anti-rat nestin (1:500, Millipore Sigma, MAB353), mouse anti-rat O4 (1:500, R&D Systems, MAB1326), rabbit anti-rat Piezo1 (1:300, Alomone, APC-087), mouse anti-rat RC2 (1:200, DSHB, RC2-C), mouse anti-rat β-tubulin III (1:500, BioLegend, 801202), and rabbit anti-rat β-tubulin III (1:500, Abcam, ab215037).

The following secondary antibodies were used: Alexa Fluor™ 488 goat anti-mouse (1:200, Invitrogen, A11001), Alexa Fluor™ 488 goat anti-mouse (1:200, Invitrogen, A21042), Alexa Fluor™ 488 goat anti-rabbit (1:200, Invitrogen, A11008), Alexa Fluor™ 568 donkey anti-goat (1:200, Invitrogen, A11057), Alexa Fluor™ 568 goat anti-mouse (1:200, Invitrogen, A11004), Alexa Fluor™ 568 goat anti-rat (1:200, Invitrogen, A11077), Alexa Fluor™ 594 goat anti-mouse (1:200, Invitrogen, A21044), Alexa Fluor™ 647 goat anti-mouse (1:200, Invitrogen, A21235), and Alexa Fluor™ 647 goat anti-rabbit (1:200, Invitrogen, A21244).

### 3.7. Image analysis

Z-stack images were projected to maximum intensity projections and thresholded for each color channel. The percentage of cells positive for a given marker or marker combination was calculated by dividing the number of cells positive in the corresponding channels by the total cell number, counted as Hoechst-positive nuclei, using ImageJ.

NPSC density in NPSC stemness maintenance experiments was calculated from the number of nuclei in the z-stack projection. Three-dimensional Sholl analysis and measurement of the number of branches on astrocyte processes, based on GFAP signal, were performed using the ImageJ plugin Simple Neurite Tracer (56) following the developers’ protocol using z-stack images. The longest protrusion distance of oligodendrocytic cells was measured from the center of the cell soma or cluster to the outermost radius of the O4 signal in z-stack projections.

To assess whether outcomes were associated with initial Ca^2+^ crosslinker concentration, marker-positive populations were replotted against the corresponding calcium sulfate concentration used for hydrogel crosslinking.

### 3.8. Flow cytometry

Cells encapsulated in hydrogels were fixed in 4% paraformaldehyde for 15 min at room temperature and permeabilized in 0.2% Triton X-100 for 15 min at room temperature. Hydrogels were transferred to microtubes, and 20 mM EDTA in PBS was added to dissociate alginate hydrogels by gentle trituration with a P1000 pipette 5 - 6 times. Then, 1% BSA in PBS was added to dilute the cell suspension, and samples were centrifuged at 500 xg for 5 min. The supernatant was removed, and cell pellets were blocked with 2.5% BSA (Sigma) in PBS for 20 min at room temperature. The suspension was centrifuged at 500 xg for 5 min. The cell pellet was resuspended in Piezo1 antibody (rabbit anti-rat Piezo1, 1:300, Alomone, APC-087) overnight at 4 °C, washed twice with 1% BSA buffer, incubated in secondary antibody (Alexa Fluor™ 488 goat anti-rabbit, 1:200, Invitrogen, A11008) for 2 h at 4 °C, and washed once with 1% BSA buffer.

Stained cells were measured for Piezo1 expression using a FACS Canto flow cytometer (BD Biosciences). Collected data were gated and analyzed using FlowJo software (BD Biosciences).

### 3.9. Mechanotransduction studies

Actomyosin-based mechanotransduction inhibition studies were performed by adding inhibitors of ROCK (Y-27632, MedChemExpress), myosin light chain kinase (ML7, MedChemExpress), and actin polymerization (cytochalasin-D, ApexBio Technology) to different media for the entire duration of culture to study their influence on differentiation outcomes. Concentrations of inhibitors were tested to maintain good cell viability during the culture period: ML7 was used at 7.5 μM for NPSC stemness maintenance, 5 μM for neuronal differentiation, 10 μM for astrocytic differentiation, 5 μM for oligodendrocytic differentiation, and 5 μM for mixed neuronal/astrocytic differentiation; Y-27632 was used at 10 μM for mixed neuronal/astrocytic differentiation; and cytochalasin-D was used at 2 μM for mixed neuronal/astrocytic differentiation. Alginate without RGD-peptide ligands was used for NPSC encapsulation to test differentiation outcomes influenced by RGD-integrin interactions.

Ion-channel-based mechanotransduction perturbation was performed by adding the Piezo1 inhibitor GsMTx4 (1 μM, Alomone) or the Piezo1 agonist Yoda1 (1 μM, Sigma) to mixed differentiation medium for the entire duration of differentiation to study their influence on differentiation outcomes. Piezo1 expression was measured on day 3 during differentiation by immunofluorescence staining followed by flow cytometry.

### 3.10. Statistical analysis

Data comparisons among more than two groups were performed using ANOVA followed by Tukey’s post-hoc tests.

## Acknowledgments

Research reported in this study was partially supported by the National Institute on Aging of the National Institutes of Health under Award Number R03AG073834. The content is solely the responsibility of the authors and does not necessarily represent the official views of the funding agencies.

## Author Contributions

S.D., J.H.K., and L.G. designed research; S.D., and J.H.K. performed experiments; S.D., J.H.K., M.W., and L.G. analyzed data; S.D., J.H.K., M.W., and L.G. wrote the paper.

