## Supporting Information for "Matrix Viscoelasticity Regulates the Stemness and Multilineage differentiation of Primary Neural Progenitor-Stem Cells in 3D"

**This PDF file includes:**

Figures S1 to S5

**Supporting Information Text**

**Figures**

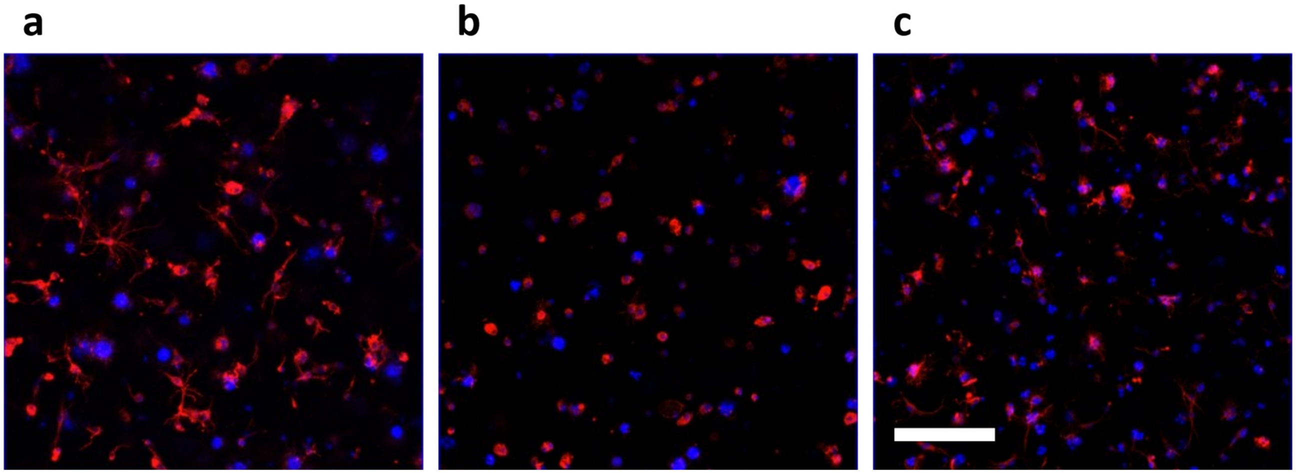


**Figure S1. NPSC spreading is reduced without adhesion ligands or with actomyosin inhibition.** (a) NPSCs cultured in fast-relaxing 7 kPa alginate hydrogels in NPSC maintenance medium. (b) NPSCs cultured in fast-relaxing 7 kPa alginate hydrogels without RGD ligands in NPSC maintenance medium. (c) NPSCs cultured in fast-relaxing 7 kPa alginate hydrogels in NPSC maintenance medium with 7.5 μM ML7 treatment. Red, nestin; blue, nuclei. Scale bar: 150 μm.

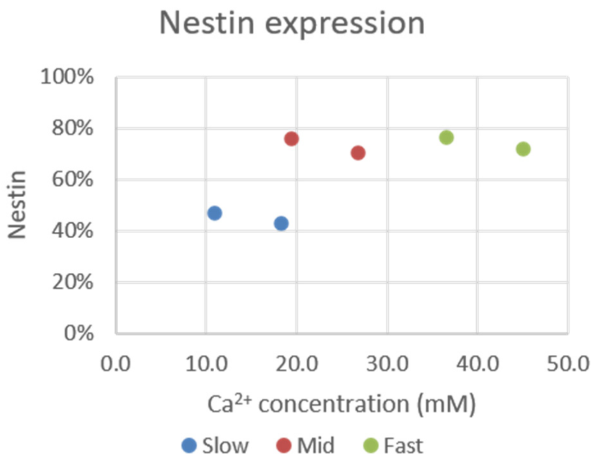


**Figure S2. Comparison of influence of initial $Ca^{2+}$ crosslinker concentration and stress relaxation on stemness maintenance.** Nestin-positive populations in NPSC maintenance medium were plotted against initial $Ca^{2+}$ crosslinker concentration, and data points were colored by stress relaxation condition.

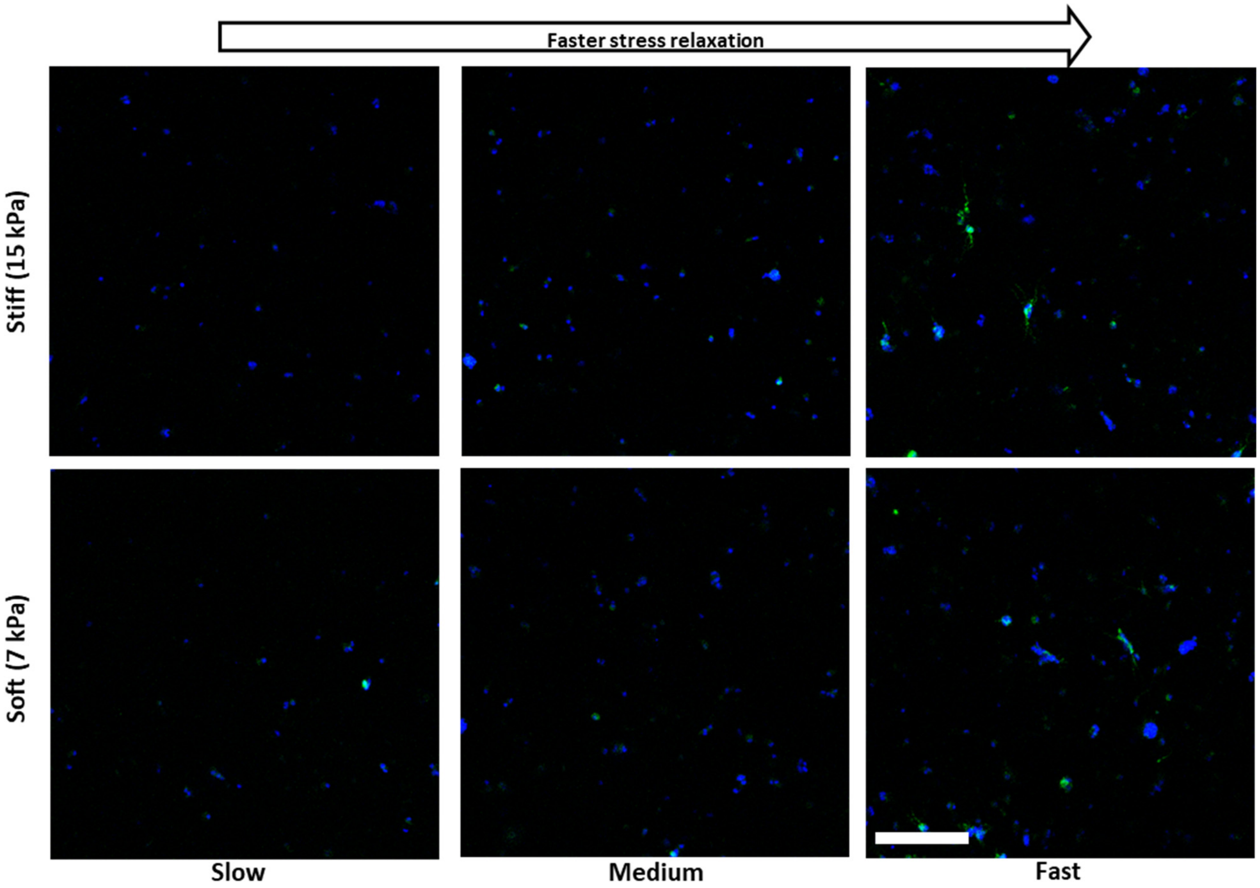


**Figure S3. MAP2ab expression after neuronal differentiation.** Representative immunocytochemistry images showing MAP2ab expression in hydrogels with different stiffness and stress relaxation properties after neuronal differentiation. Green, MAP2ab; blue, nuclei. Scale bar: 150 μm.

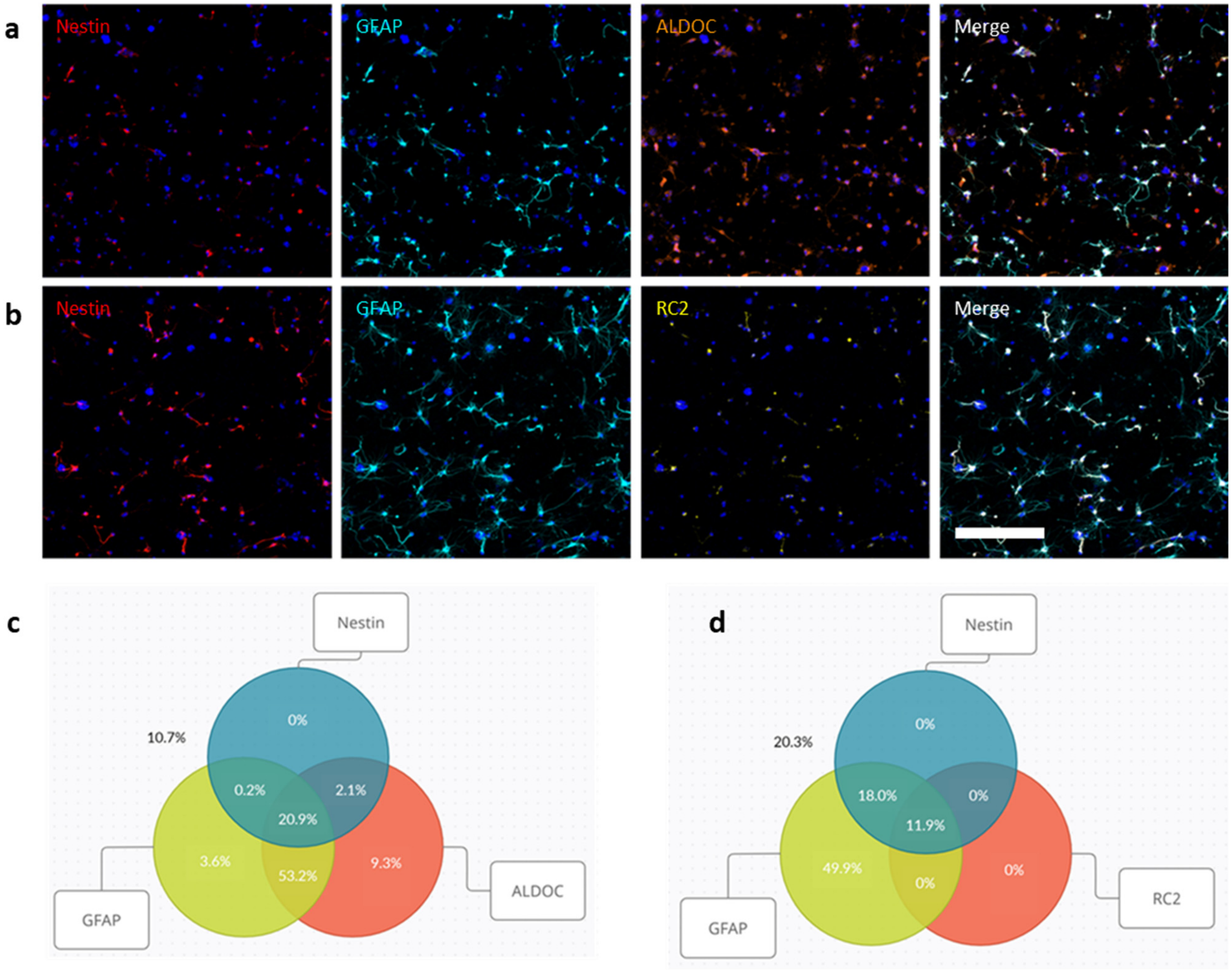


**Figure S4. Characterization of nestin$^{-}$GFAP$^{+}$ cells after astrocytic differentiation of NPSCs in fast-relaxing 7 kPa hydrogels.** (a) Nestin$^{-}$GFAP$^{+}$ cells express ALDOC. Red, nestin; cyan, GFAP; orange, ALDOC. (b) Nestin$^{-}$GFAP$^{+}$ cells lack RC2 expression. Red, nestin; cyan, GFAP; yellow, RC2. Quantitative results are shown in (c) and (d), respectively. Scale bar: 150 μm.

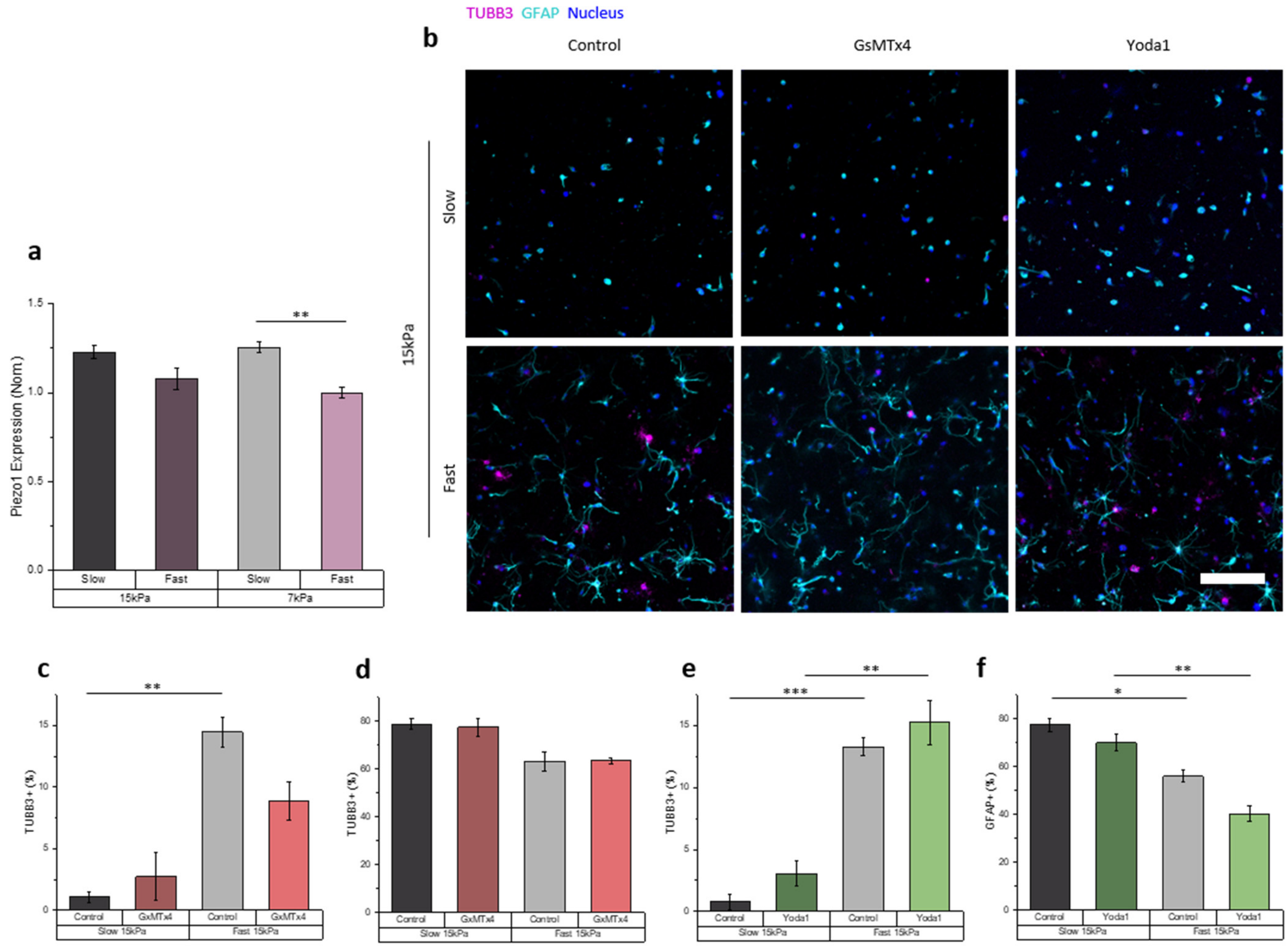


**Figure S5. Piezo1 expression and activity modulation in mixed differentiation culture condition.** (a) Piezo1 expression quantified from fluorescence cytometry shown in normalized mean fluorescence intensity (N=5). (b) Immunocytochemistry in environments of different stiffness and stress-relaxation shows that TUBB3 and GFAP expression and cell spreading vary with GsMTx4 or Yoda1. Magenta – TUBB3; cyan – GFAP; blue – nucleus. (c,d) Quantification of TUBB3 or GFAP positive cell population in addition of GsMTx4 inhibitor (N=3). (e,f) Quantification of TUBB3 or GFAP positive cell population in addition of Yoda1 agonist (N=3). * indicates $p<0.05$, ** indicates $p<0.01$, *** indicates $p<0.001$ (ANOVA with Tukey post-hoc test). Data are shown in means with error bars indicating ±s.e.m. Scale bar, 150 μm.
